# Chromosome-scale assembly of the genome of *Salix dunnii* reveals a male-heterogametic sex determination system on chromosome 7

**DOI:** 10.1101/2020.10.09.333229

**Authors:** Li He, Kai-Hua Jia, Ren-Gang Zhang, Yuan Wang, Tian-Le Shi, Zhi-Chao Li, Si-Wen Zeng, Xin-Jie Cai, Natascha Dorothea Wagner, Elvira Hörandl, Aline Muyle, Ke Yang, Deborah Charlesworth, Jian-Feng Mao

## Abstract

Sex determination systems in plants can involve either female or male heterogamety (ZW or XY, respectively). Here we used Illumina short reads, Oxford Nanopore Technologies (ONT) long reads, and Hi-C reads to assemble the first chromosome-scale genome of a female willow tree (*Salix dunnii*), and to predict genes using transcriptome sequences and available databases. The final genome sequence of 328 Mb in total was assembled in 29 contigs, and includes 31,501 genes. We inferred a male heterogametic sex determining factor on chromosome 7, suggesting that, unlike the female heterogamety of most species in the genus *Salix*, male heterogamety evolved in the subgenus *Salix*. The *S. dunnii* X-linked region occupies about 3.21 Mb of chromosome 7, and is probably in a pericentromeric region. Our data suggest that this region is enriched for transposable element insertions, and about one third of its 124 protein-coding genes were gained via duplications from other genome regions. We detect purifying selection on the genes that were ancestrally present in the region, though some have been lost. Transcriptome data from female and male individuals show more male- than female-biased genes in catkin and leaf tissues, and indicate enrichment for male-biased genes in the pseudo-autosomal regions. Our study provides valuable genomic resources for studying sex chromosome evolution in Salicaceae family.

## Introduction

Dioecious plants are found in approximately 5-6 % of flowering plant species (Charlesworth 1985; Renner 2014), and genetic sex determination systems have evolved repeatedly among flowering plants, and independently in different lineages, and include a range of evolutionary stages, some species having pronounced morphological differences between their sex chromosomes (heteromorphism), while others have homomorphic sex chromosomes (reviewed by Westergaard 1958; Ming *et al*. 2011). Recent progress has included identifying sex-linked regions in several plants with homomorphic sex chromosomes, and some of these have been found to be small parts of the chromosome pairs, allowing sex determining genes to be identified (e.g. Harkess *et al*. 2017; Akagi *et al*. 2019; Harkess *et al*. 2020; Zhou *et al*. 2020; Müller *et al*. 2020); the genes are often involved in hormone response pathways, mainly associated with cytokinin and ethylene response pathways (reviewed by Feng *et al*. 2020). XX/XY (male heterogametic) and ZW/ZZ (female heterogametic) sex determination systems have been found in close relatives (Balounova *et al*. 2019; Martin *et al*. 2019). However, the extent to which related dioecious plants share the same sex-determining systems, or evolved dioecy independently, is still not well understood.

After recombination stops between an evolving sex chromosome pair, or part of the pair, forming a fully sex-linked region, repetitive sequences and transposable elements are predicted to accumulate rapidly (reviewed in Bergero & Charlesworth 2009). The expected accumulation has been detected in both Y- and W-linked regions of several plants with heteromorphic sex chromosome pairs (reviewed by Hobza *et al*. 2015). Repeat accumulation is also expected in X- and Z-linked regions. Although accumulation is expected to occur to a much smaller extent, it has been detected in the X of *Carica papaya* (Gschwend *et al*. 2012; Wang *et al*. 2012a), and, in a willow species, *S. purpurea*, both the Z and the W show almost the same large enrichment (Zhou *et al*. 2020). The accumulation of repeats reduces gene densities, compared with autosomal or pseudoautosomal regions (PARs), and this has been observed in *Silene latifolia*, again affecting both sex chromosomes (Blavet *et al*. 2015), and in the *S. purpurea* Z and W (Zhou *et al*. 2020).

The accumulation of repetitive sequences is a predicted consequence of recombination suppression reducing the efficacy of selection in Y and W-linked regions compared to those carried on X and Z chromosomes, which also predicts that deleterious mutations will accumulate, causing Y and W chromosome genetic degeneration (reviewed by Charlesworth *et al*. 1994, Ellegren 2011 and Wang *et al*. 2012a). The chromosome that recombines in the homogametic sex (the X or Z) remains undegenerated and maintains the ancestral gene content of its progenitor chromosome, and purifying selection can act to maintain genes’ functions (Wilson & Makova 2009). However, genes on these chromosomes are also predicted to evolve differently from autosomal genes. Compared with purifying selection acting on autosomal genes, hemizygosity of genes in degenerated regions increases the effectiveness of selection against X- or Z-linked deleterious mutations (unless they are not expressed in the heterogametic sex, see Vicoso & Charlesworth 2006). Positive selection may also act on X/Z-linked genes, and will be particularly effective in causing spread of X-linked male-beneficial mutations (or Z in female-beneficial ones in ZW systems), because mutations are hemizygous in the heterogametic sex (Vicoso & Charlesworth 2006). When comparing coding sequences between different species, X and Z-linked genes may therefore have either higher *K*a/*K*s (non-synonymous substitution per non-synonymous site/synonymous substitution per synonymous site) ratios than autosomal genes, or lower ratios if purifying selection against deleterious mutations is more important (Vicoso & Charlesworth 2006). Furthermore, X/Z-linked regions may, over time, gain genes with beneficial effects in one sex, but deleterious effects in the other (sexually antagonistic effects, see Rice 1984; Arunkumar *et al*. 2009; Meisel *et al*. 2012).

Here, we studied a member of the Salicaceae, a plant family that may include multiple evolutionary origins of genetic sex-determination. The family *sensu lato* (*s*.*l*.) includes more than 50 genera and 1,000 species, usually dioecious or monoecious (rarely hermaphroditic) (Chase *et al*. 2002; Cronk *et al*. 2015). Roughly half of the species are in two closely related genera of woody trees and shrubs, *Populus* and *Salix*, whose species are almost all dioecious (Fang *et al*. 1999; Argus 2010), which might suggest that dioecy is the ancestral state. However, studies over the past 6 years, summarized in Table 1, show that the sex-linked regions are located in different genome regions in different species, and that both genera include species whose sex-determining regions appear to be in the early stages in the evolution.

**Table 1.**
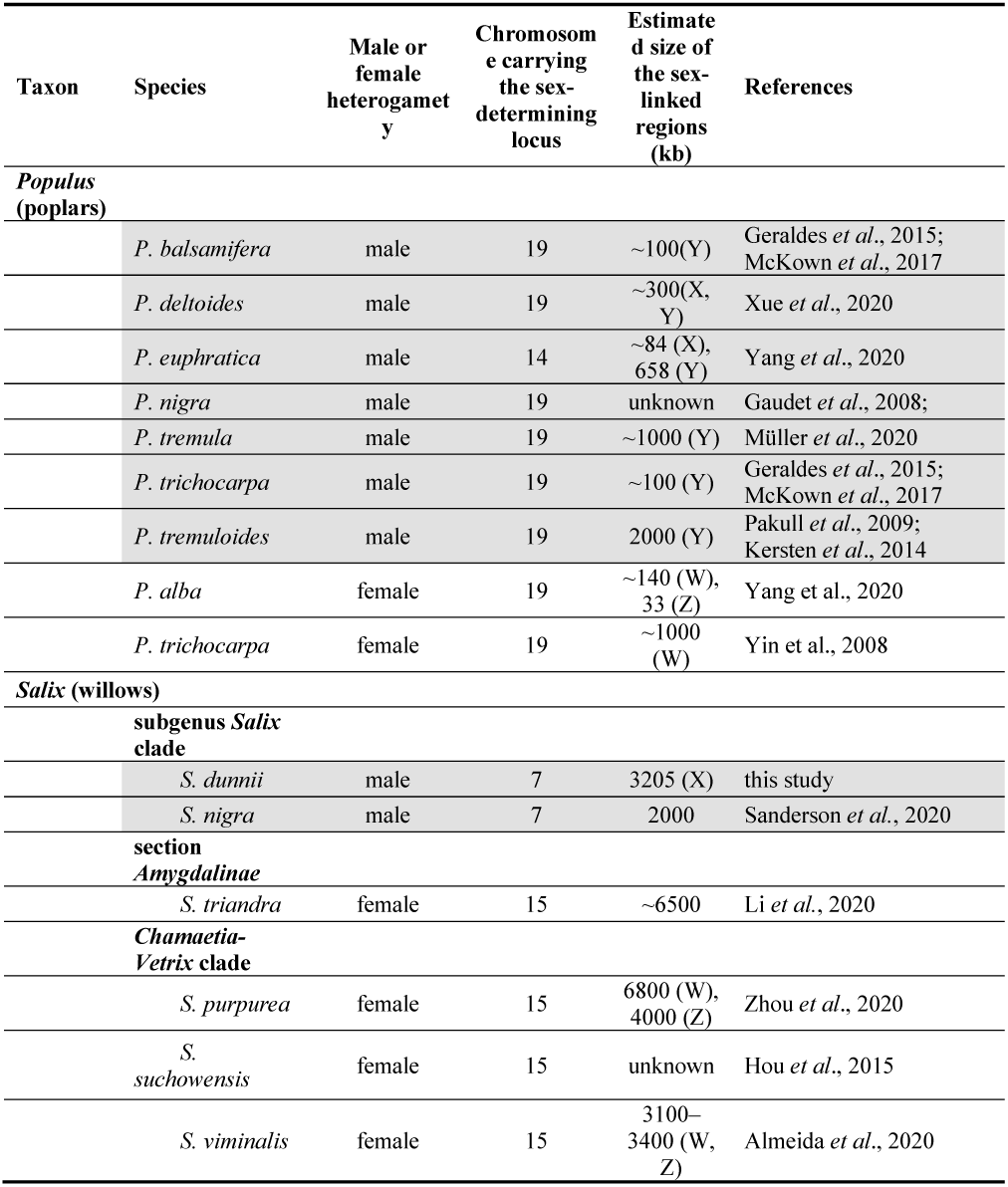
Summarizing current information about sex-linked regions in *Populus* and *Salix*.

*Populus* species usually have XX/XY systems and sex determining regions (SDR) on chromosome 14 or 19, though a few species have ZW/ZZ systems with the SDR also on chromosome 19. Until recently, all willows investigated were from one *Salix* clade, *Chamaetia-Vetrix* (Lauron-Moreau *et al*. 2015; Wu *et al*. 2015), and all were found to have female heterogamety and SDRs on chromosome 15 (Table 1), as does the close relative *S. triandra* (section *Amygdalinae*), but, as the table shows, a recent study suggested an XX/XY system on chromosome 7 in *S. nigra*, the only species so far studied from subgenus *Salix* clade (sensu Wu *et al*. 2015). This evidence for changes in the location of the sex-linked regions, and for differences in the heterozygous sex, make the family Salicaceae interesting for studying the evolution of sex chromosomes, and in particular sex chromosome turnover.

To understand the evolutionary events involved in these differences, high-quality genome sequences are needed, leading, potentially, to discovery of the sex-determining gene(s), which can reveal whether the same gene is involved in species with the same heterogamety (perhaps even across different genera), or whether different lineages have independently evolved sex-determining systems. Recent studies in *Populus* identified a gene in the *Arabidopsis thaliana* Type A response regulator family (resembling ARABIDOPSIS RESPONSE REGULATOR 17, and therefore named *ARR17*), within the sex-linked region on chromosome 19 of both *P. tremula* and *P. deltoides*, which has been shown to be involved in sex-determination in *P. tremula* (Müller *et al*. 2020; Xue *et al*. 2020). In two species of the *Salix Chamaetia-Vetrix* clade (*S. purpurea* and *S. viminalis*), a similar gene has been detected in the W-linked region of chromosome 15, and a partial and non-functional copy was found in the Z-linked region of the *S. purpurea* chromosome 15 (Almeida *et al*. 2020; Yang *et al*. 2020; Zhou *et al*. 2020). However, these are moderately extensive regions (see Table 1) and, given that this gene family includes several members, the finding of such copies is not strong evidence that such a gene is involved in sex-determination in willows. Studying other willow species might confirm presence of such a gene in all willow SDRs, or might instead find that some species’ SDRs include no such gene. Species with different heterogamety are of special interest, because these seem unlikely to involve the same gene.

Although *Salix* is the largest genus in the family Salicaceae *sensu lato*, with ∼450 species (reviewed in He *et al*. 2020), fewer *Salix* than *Populus* genomes have been assembled, and assemblies include only the cushion shrub *S. brachista* and the shrub willows *S. purpurea, S. suchowensis*, and *S. viminalis* (Chen *et al*. 2019; Almeida *et al*. 2020; Wei *et al*. 2020; Zhou *et al*. 2020). Shrub stature is a derived character, and the tree habit is ancestral (Skvortsov 1999), and is usual in poplars.

Here, we describe studies in *S. dunnii*, a riparian willow tree of the subgenus *Salix* clade (sensu Wu *et al*. 2015), found in subtropical areas of China that can grow up to 10 meters (Fang *et al*. 1999). Our study has three aims. First, to develop a high quality, chromosome level assembly of the *S. dunnii* genome, which has not previously been sequenced. Second, to re-sequence samples of both sexes from natural populations to test whether this subgenus *Salix* species has an XX/XY system, and, if so, whether it is on chromosome 7, as in *S. nigra*, suggesting a possible independent evolutionary origin from the ZW systems in other *Salix* clades. Third, to study the evolution of the X-linked region. Several interesting questions include (i) whether recombination in the region has changed since it became an X-linked region (versus a sex-determining region having evolved within an already non-recombining region), (ii) whether the genes in the region are orthologs of those in the homologous region of related species (versus genes having been gained by movements from other genome regions), and (iii) whether the X-linked region genes differ in expression between the sexes, and/or (iv) have undergone adaptive changes more often than other genes.

## Materials and Methods

### Plant material

We collected young leaves from a female *S. dunnii* plant (FAFU-HL-1) for genome sequencing. Silica-gel dried leaves were used to estimate ploidy. Young leaf, catkin, stem, and root samples for transcriptome sequencing were collected from female FAFU-HL-1, and catkins and leaves from two other female and three male plants. We sampled 38 individuals from two wild populations of *S. dunnii* for resequencing. The plant material was frozen in liquid nitrogen and stored at −80°C until total genomic DNA or RNA extraction. For sequencing involving Oxford Nanopore Technologies (ONT) and Hi-C, fresh leaf material was used. Table S1 gives detailed information about all the samples.

### Ploidy determination

The ploidy of FAFU-HL-1 was measured by flow cytometry (FCM), using a species of known ploidy (*Salix integra*; 2x = 2n = 38, Wagner *et al*. 2020) as an external standard. The assay followed the FCM protocol of Doležel *et al*. (2007) (see Supplementary Note 1).

### Genome sequencing

For Illumina PCR-free sequencing, total genomic DNA of FAFU-HL-1 was extracted using a Qiagen DNeasy Plant Mini kit following the manufacturer’s instructions (Qiagen, Valencia, CA). For ONT sequencing, phenol-chloroform was used to extract DNA. PCR-free sequencing libraries were generated using Illumina TruSeq DNA PCR-Free Library Preparation Kit (Illumina, USA) following the manufacturer’s recommendations. After quality assessment on an Agilent Bioanalyzer 2100 system, the libraries were sequenced on an Illumina platform (NovaSeq 6000) by Beijing Novogene Bioinformatics Technology Co., Ltd. (hereafter referred to as Novogene). ONT libraries were prepared following the Oxford Nanopore 1D Genomic DNA (SQKLSK109)-PromethION ligation protocol, and sequenced by Novogene.

### Hi-C library preparation and sequencing

The Hi-C library was prepared following a standard procedure (Wang *et al*. 2020). In brief, fresh leaves from FAFU-HL-1 were fixed with a 1% formaldehyde solution in MS buffer. Subsequently, cross-linked DNA was isolated from nuclei. The DPNII restriction enzyme was then used to digest the DNA, and the digested fragments were labeled with biotin, purified, and ligated before sequencing. Hi-C libraries were controlled for quality and sequenced on an Illumina Hiseq X Ten platform by Novogene.

### RNA extraction and library preparation

Total RNA was extracted from young leaves, female catkins, stems, and roots of FAFU-HL-1 using the Plant RNA Purification Reagent (Invitrogen) according to the manufacturer’s instructions. Genomic DNA was removed using DNase I (TaKara). An RNA-seq transcriptome library was prepared using the TruSeq™ RNA sample preparation Kit from Illumina (San Diego, CA) and sequencing was performed on an Illumina Novaseq 6000 by the Shanghai Majorbio Bio-pharm Biotechnology Co., Ltd., China (hereafter referred to as Majorbio).

### Genome size estimation

The genome size was estimated by 17-k-mer analysis based on PCR-free Illumina short reads to be ∼376 Mb. Briefly, k-mer were counted using JELLYFISH (Marçais *et al*. 2011), and the numbers used to estimate the genome size and repeat content using findGSE (Sun *et al*. 2018). The proportion of sites in this individual that are heterozygous was estimated using GenomeScope (Vurture *et al*. 2017).

### Genome assembly

SMARTdenovo (Ruan 2016) and wtdbg2 (Ruan & Li 2020) were used to create a *de novo* assembly based on ONT reads, using the following options: -c l to generate a consensus sequence, -J 5000 to remove sequences <5 kb, and -k 20 to use 20-mers. We then selected the assembly with the highest N50 value and a genome size close to the estimated one, which was assembled by SMARTdenovo with Canu correction (Koren *et al*. 2017) (Table S2). Since ONT reads contain systematic errors in regions with homo-polymers, we mapped Illumina short reads to the genome and polished using Pilon (Walker *et al*. 2014). The Illumina short reads were filtered using fastp (Chen *et al*. 2018) to remove adapters and low base quality sequences before mapping.

### Scaffolding with Hi-C data

We filtered Hi-C reads using fastp (Chen *et al*. 2018), then mapped the clean reads to the assembled genome with Juicer (Durand *et al*. 2016), and finally assembled them using the 3d-DNA pipeline (Dudchenko *et al*. 2017). Using Juicebox (Durand *et al*. 2016), we manually cut the boundaries of chromosomes. In order to decrease the influence of inter-chromosome interactions and improve the chromosome-scale assembly, we separately re-scaffolded each chromosome with 3d-DNA, and further corrected mis-joins, order, and orientation of a candidate chromosome-length assembly using Juicebox. Finally, we anchored the contigs to 19 chromosomes. The *Rabl* configuration (Dong & Jiang 1998; Prieto *et al*. 2004) is not obvious enough to predict the possible centromere positions in chromosome 7 of *S. dunnii* (Figure S1), but we employed Minimap2 (Li 2018) with parameters “-x asm20”, in order to identify the region with highest repeat sequence densities in the genome, which may represent the centromere.

### Optimizing the genome assembly

To further improve the genome assembly, LR_Gapcloser (Xu *et al*. 2019a) was employed twice for gap closing with ONT reads. We also used NextPolish (Hu *et al*. 2020) to polish the assembly, with three iterations with Illumina short reads to improve base accuracy. We subsequently removed contigs with identity of more than 90% and overlap of more than 80 %, which were regarded as redundant sequences, using Redundans (Pryszcz *et al*. 2016), Overall, we removed a total of 8.62 Mb (40 contigs) redundant sequences. Redundant sequences were mainly from the same regions of homeologous chromosomes (Pryszcz *et al*. 2016). To identify and remove contaminating sequences from other species, we used the contigs to blast against the NCBI-NT database, and found no contaminated contigs.

### Characterization of repetitive sequences

Repeat elements were identified and classified using RepeatModeler (http://www.repeatmasker.org/) to produce a repeat library. Then RepeatMasker was used to identify repeated regions in the genome, based on the library. The repeat-masked genome was subsequently used in gene annotation.

### Annotation of full-length LTR-RTs and estimation of insertion times

We annotated full-length LTR-RTs in our assembly and estimated their insertion times as described in Xu *et al*. (2019b). Briefly, LTRharvest (Ellinghaus *et al*. 2008) and LTRdigest (Steinbiss *et al*. 2009) were used to *de novo* predict full-length LTR-RTs in our assembly. LTR-RTs were then extracted and compared with *Gag-Pol* protein sequences within the REXdb database (Neumann *et al*. 2019). To estimate their insertion times, the LTRs of individual transposon insertions were aligned using MAFFT (Katoh & Standley 2013), and divergence between the 5’and 3’-LTR was estimated (Sanmiguel 1998; Ma & Bennetzen 2004). The divergence values were corrected for saturation by Kimura’s two-parameter method (Kimura 1980), and insertion times were estimated from the values, assuming a mutation rate of 2.5×10 ^−9^ substitutions year^−1^ per site (Ingvarsson 2008).

### Transcriptome assembly and gene annotation

The genome was annotated by combining evidence from transcriptome, *ab initio* prediction, and protein homology based on prediction. PASA (Program to Assemble Spliced Alignment, Haas *et al*. 2003) was used to obtain high-quality loci based on transcriptome data. We randomly selected half of these loci as a training dataset to train the AUGUSTUS (Stanke *et al*. 2008) gene modeller, and the other half as the test dataset, and conducted five replicates of optimization. The high-quality loci data set was also used to train SNAP (Korf 2004). A total of 103,540 protein sequences were obtained from *Arabidopsis thaliana, P. trichocarpa, S. purpurea* and *S. suchowensis* and used as reference proteins for homology-based gene annotation. Gene annotation was then performed with the MAKER pipeline (Cantarel *et al*. 2008) (Detail process presented in Supplementary Note 2).

To annotate tRNA and rRNA sequences, we used tRNAScan-SE (Lowe & Eddy 1997) and RNAMMER (Lagesen *et al*. 2007), respectively, and other ncRNAs were identified by querying against the Rfam database (Nawrocki *et al*. 2015).

For protein functional annotation, the annotated genes were aligned to proteins in Uniprot database (including the SWISS-PROT and TrEMBL databases, https://www.uniprot.org/), NR (https://www.ncbi.nlm.nih.gov/), Pfam and eggNOG (Powell *et al*. 2014) databases using BLAT (E value <10^−5^) (Kent 2002). Motifs and functional domains were identified by searching against various domain libraries (ProDom, PRINTS, Pfam, SMART, PANTHER and PROSITE) using InterProScan (Jones *et al*. 2014). Annotations were also assigned to GO (http://geneontology.org/) and KEGG (https://www.genome.jp/kegg/pathway.html) metabolic pathways to obtain more functional information.

To identify pseudogenes, the proteins were aligned against the genome sequence using tBLASTn with parameter settings of “-m 8 -e 1e-5”. PseudoPipe with default parameter settings was then used to detect pseudogenes in the whole genome (Zhang *et al*. 2006).

### Comparative phylogeny analysis across willows

We performed a comparative genomic investigation of the available willow genomes (*Salix dunnii, S. brachista, S. purpurea, S. suchowensis*, and *S. viminalis*), used *Populus trichocarpa* as an outgroup (Table S3). OrthoFinder2 (Emms & Kelly 2018) was used to identify groups of orthologous genes. A maximum likelihood (ML) phylogenetic tree was constructed using IQ-TREE (Nguyen *et al*. 2014) based on single-copy orthologs extracted from orthogroups. The CDS (Coding DNA Sequence) of the single-copy orthologous genes identified were aligned with MAFFT (Katoh & Standley 2013), and then trimmed with trimAI (Capella-Gutiérrez *et al*. 2009). Finally, MCMCTree in the PAML package (Yang 2007) was used to estimate the divergence time. For more details, see Supplementary Note 3.

We performed collinearity analysis of *P. trichocarpa* and the five willows, and self-comparison of each species, using MCScanX with the default parameters (Wang *et al*. 2012b). KaKs_Calculator (Wang *et al*. 2010) was used to calculate *K*a (the number of substitutions per nonsynonymous site), *K*s (substitutions per synonymous site), and *K*a/*K*s values, based on orthologous pairs, using the Yang-Nielsen (YN) model (Zhang & Yu 2006).

### Whole-genome resequencing and SNP calling

Total genomic DNA for all 38 samples (Table S1) was extracted with the Qiagen DNeasy Plant Mini Kit (Qiagen, Valencia, CA) following the manufacturer’s instructions. Whole-genome resequencing using paired-end libraries was performed on Illumina NovaSeq 6000 by Majorbio. The sequenced reads were filtered and trimmed by fastp (Chen *et al*. 2018). The filtered reads were then aligned to the assembled genome using the BWA-MEM algorithm from BWA (Li & Durbin 2009; Li 2013). SAMtools (Li *et al*. 2009) was used to extract primary alignments, sort, and merge the mapped data. Sambamba (Tarasov *et al*. 2015) was used to mark potential duplications in the PCR amplification step of library preparation. Finally, FreeBayes (Garrison & Marth 2012) was employed for SNP calling, yielding 10,985,651 SNPs. VCFtools software (Danecek *et al*. 2011) was used to select high-quality SNPs based on the calling results: we (1) excluded all genotypes with a quality below 20, (2) included only genotypes with coverage depth at least 5 and not more than 200, (3) retained only bi-allelic SNPs, (4) removed SNPs with missing information rate > 20% and minor allele frequency < 5%. This yielded 4,370,362 high-quality SNPs for analysis.

### Identification of sex determination systems

We used our high-quality SNPs in a standard case-control genome-wide association study (GWAS) between allele frequencies and sex phenotype using PLINK (Purcell *et al*. 2007). SNPs with α < 0.05 after Bonferroni correction for multiple testing were considered significantly associated with sex.

The chromosome quotient (CQ) method (Hall *et al*. 2013) was employed to further test whether *S. dunnii* has a female or male heterogametic system. The CQ is the normalized ratio of female to male alignments to a given reference sequence, using the stringent criterion that the entire read must align with zero mismatches. To avoid bias due to different numbers of males and females, we used only 18 individuals of each sex (Table S1). We filtered the reads with fastp, and made combined female and male read datasets. The CQ-calculate.pl software (https://sourceforge.net/projects/cqcalculate/files/CQ-calculate.pl/download) was used to calculate the CQ for each 50 kb nonoverlapping window of the *S. dunnii* genome. For male heterogamety, we expect a CQ value close to 2 in windows in the X-linked region (denoted below by X-LR), given a female genome sequence, whereas, for female heterogamety we expect CQ ≈ 0.5 for Z-linked windows, and close to zero for W-linked windows.

Population genetic statistics, including nucleotide diversity per base pair (π) and observed heterozygote frequencies (*H*_obs_) were calculated for female and male populations using VCFtools (Danecek *et al*. 2011) or the “populations” module in Stacks (Catchen *et al*. 2011). Weighted *F*_ST_ values between the sexes were calculated using the Weir & Cockerham (1984) estimator with 100 kb windows and 5 kb steps. A Changepoint package (Killick & Eckley 2014) was used to assess significance of differences in the mean and variance of the *F*_ST_ values between the sexes of chromosome 7 windows, using function cpt.meanvar, algorithm PELT and penalty CROPS. PopLDdecay (Zhang *et al*. 2019) was used to estimate linkage disequilibrium (LD) based on unphased data, for the whole genome and the X-LR, with parameters “-MaxDist 300 -MAF 0.05 -Miss 0.2”.

### Gene content of chromosome 7 of *Salix dunnii*

MCscan (Python version) (Tang *et al*. 2008) was used to analyze chromosome collinearity between the protein-coding sequences detected in the whole genomes of *S. dunnii, S. purpurea* and *P. trichocarpa*. The “--cscore=.99” was used to obtain reciprocal best hit (RBH) orthologs for synteny analysis.

To identify homologous gene pairs shared by chromosome 7 and the autosomes of *S. dunnii*, and those shared with chromosome 7 of *P. trichocarpa*, and *S. purpurea* (using the genome data in Table S3), we did reciprocal blasts of all primary annotated peptide sequences with “blastp -evalue 1e-5 -max_target_seqs 1”. For genes with multiple isoforms, only the longest one was used. Furthermore, homologs of *S. dunnii* chromosome 7 genes *in Arabidopsis thaliana* were identified with same parameters.

Because the *A. thaliana* ARR17 gene (AT3G56380.1, https://www.arabidopsis.org/) has been proposed to be a sex-determining gene in *Salix* (see Introduction), we also blasted its sequence against our assembled genome with “tblastn -max_target_seqs 5 -evalue 1e-5” to identify possible homologous intact or pseudogene copies.

### Molecular evolution of chromosome 7 homologs of willow and poplar

To test whether X-linked genes in our female genome sequence evolve differently from other genes, we aligned homologs of chromosome 7 sequences identified by blastp, and estimated the value of *K*a and *K*s between *S. dunnii* and *P. trichocarpa*, and between *S. dunnii* and *S. purpurea*. To obtain estimates for an autosome for the same species pairs, we repeated this analysis for chromosome 6 (this is the longest chromosome, apart from chromosome 16, which has a different arrangement in poplars and willows, see Results, Table S4). ParaAT (Zhang *et al*. 2012) and Clustalw2 (Larkin *et al*. 2007) were used to align the sequences, and the yn00 package of PAML (Yang 2007) was used to calculate the *K*a and *K*s values for each homologous pair.

### Gene expression

We used Seqprep (https://github.com/jstjohn/SeqPrep) and Sickle (https://github.com/najoshi/sickle) to trim and filter the raw data from 12 tissue samples (catkins and leaves from each of three female and male individuals) (Table S1).

Clean reads were separately mapped to our assembled genome for each sample using STAR (Dobin *et al*. 2013) with parameters “--sjdbOverhang 150, --genomeSAindexNbases 13”. The featureCounts program (Liao *et al*. 2014) was employed to merge different transcripts to a consensus transcriptome and calculate counts separately for each sex and tissue. Then we converted the read counts to TPM (Transcripts per million reads), after filtering out unexpressed genes (counts=0 in all samples, excluding non-mRNA). 28,177 (89.45%) genes were used for subsequent analyses. The DEseq2 package (Love *et al*. 2014) was used to detect genes differentially expressed in the different sample groups. The DESeq default was used to test differential expression using negative binomial generalized linear models and estimation of dispersion and logarithmic fold changes incorporating data-driven prior distributions, to yield log_2_FoldChanges values and p values adjusted for multiple tests (adjusted p value < 0.05, | log_2_FoldChange | > 1).

## Results

### Genome assembly

*k*-mer analysis of our sequenced genome of a female *S. dunnii* plant indicated that the frequency of heterozygous sites in this diploid individual is low (0.79%) (Figure S2, Figure S3; Table S1). We generated 72Gb (∼180×) of ONT long reads, 60 Gb (∼150×) Illumina reads, and 55 Gb (∼140×) of Hi-C reads (Table S5, Table S6). After applying several different assembly strategies, we selected the one with the ‘best’ contiguity metrics (SMARTdenovo with Canu correction, Table S2). Polishing/correcting using Illumina short reads of the same individual yielded a 333 Mb genome assembly in 100 contigs (contig N50 = 10.1 Mb) (Table S2).

With the help of Hi-C scaffolding, we achieved a final chromosome-scale assembly of 328 Mb of 29 contigs (contig N50 = 16.66 Mb), about 325.35 Mb (99.17%) of which is anchored to 19 pseudochromosomes (scaffold N50 = 17.28 Mb) (Figure 1a and Figure S4; Table 2 and Table S4), corresponding to the haploid chromosome number of the species. The mitochondrial and chloroplast genomes were assembled into circular DNA molecules of 711,422 bp and 155,620 bp, respectively (Figure S5, Figure S6). About 98.4% of our Illumina short reads were successfully mapped back to the genome assembly, and about 99.5% of the assembly was covered by at least 20× reads. Similarly, 98.9% of ONT reads mapped back to the genome assembly and 99.9% were covered by at least 20× reads. The assembly’s LTR Assembly Index (LAI) score was 12.7, indicating that our assembly reached a high enough quality to achieve the rank of “reference” (Ou *et al*. 2018). BUSCO (Simão *et al*. 2015) analysis identified 1,392 (96.6%) of the 1,440 highly conserved core proteins in the Embryophyta database, of which 1,239 (86.0%) were single-copy genes and 153 (10.6%) were duplicate genes. A further 33 (2.3%) had fragmented matches to other conserved genes, and 37 (2.6%) were missing.

**Table 2.**
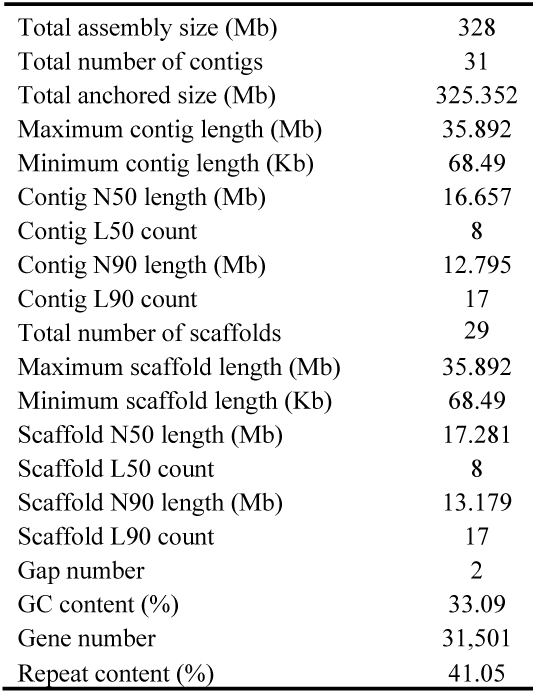
Statistics of the *Salix dunnii* genome assembly.

**Fig. 1.**
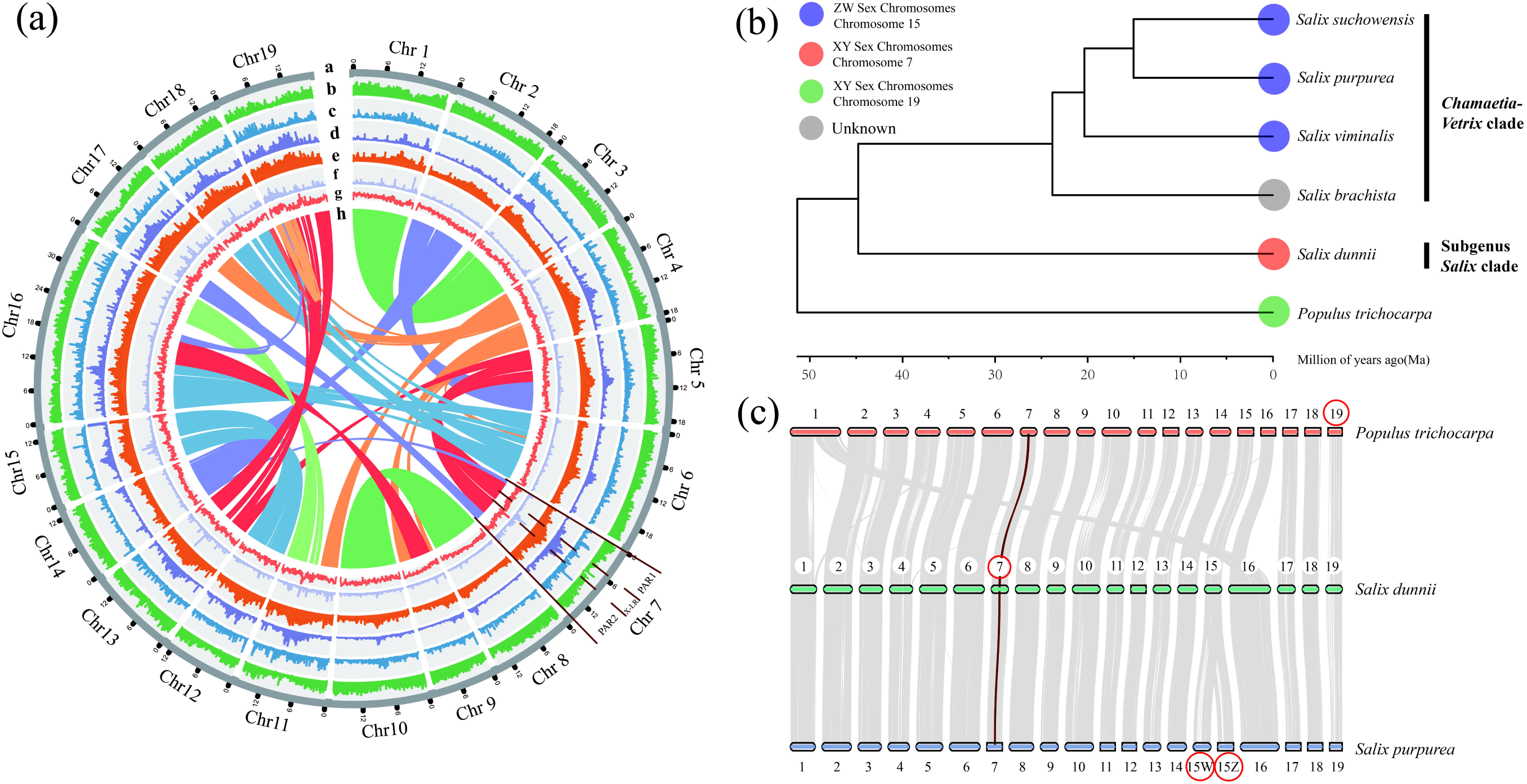
Genome structure and evolution of *S. dunnii*. **a**, Circos plot showing: (a) the chromosome lengths in Mb, (b) gene density, (c) LTR-Copia density, (d) LTR-Gypsy density, (e) total repeats, (f) density of pseudogenes, (g) GC (guanine-cytosine) content, (h) Syntenic blocks. **b**, Inferred phylogenetic tree of *S. dunnii, S. viminalis, S. brachista, S. purpurea, S. suchowenssi* and the outgroup *P. trichocarpa*, with divergence times. The root age of the tree was calibrated to 48–52 Ma following Chen *et al*. (2019) and the crown age of the *Chamaetia-Vetrix* clade (here including *S. brachista, S. purpurea, S. suchowensis*, and *S. viminalis*) was calibrated to 23–25 Ma according to Wu *et al*. (2015). **c**, Macrosynteny between genomic regions of *P. trichocarpa, S. dunnii*, and *S. purpurea*. The dark orange line shows the syntenic regions between the *S. dunnii* X-linked region of chromosome 7, and the homologous regions in the same chromosomes of *S. purpurea* and *P trichocarpa*. Red circles show the chromosomes carrying sex linked region.

### Annotation of genes and repeats

134.68 Mb (41.0%) of the assembled genome consisted of repetitive regions (Table 2), close to the 41.4 % predicted by findGSE (Sun *et al*. 2018). Long terminal repeat retrotransposons (LTR-RTs) were the most abundant annotations, forming up to 19.1% of the genome, with *Gypsy* and *Copia* class I retrotransposon (RT) transposable elements (TEs) accounting for 13% and 5.85% of the genome, respectively (Table S7). All genomes so far studied in *Salix* species have considerable proportions of transposable element sequences, but the higher proportions of *Gypsy* elements in *S. dunnii* (Table S7) (Chen *et al*. 2019) suggested considerable expansion in this species. Most full-length LTR-RTs appear to have inserted at different times within the last 30 million years rather than in a recent burst (Figures S7, S8, and S9; Table S8).

Using a comprehensive strategy combining evidence-based and *ab initio* gene prediction (see Methods), we then annotated the repeat-masked genome. We identified a total of 31,501 gene models, including 30,200 protein-coding genes, 650 transfer RNAs (tRNAs), 156 ribosomal RNAs (rRNA) and 495 unclassifiable non-coding RNAs (ncRNAs) (Table 2; Table S9). The average *S. dunnii* gene is 4,095.84 bp long and contains 6.07 exons (Table S10). Most of the predicted protein-coding genes (94.68%) matched a predicted protein in a public database (Table S11). Among the protein-coding genes, 2,053 transcription factor (TF) genes were predicted and classified into 58 gene families (Table S12, Table S13).

### Comparative genomics and whole genome duplication events

We compared the *S. dunnii* genome to those of four published willow genomes and *Populus trichocarpa* as an outgroup, using 5,950 single-copy genes to construct a phylogenetic tree of the species’ relationships (Figure 1b). Consistent with published topologies (Wu *et al*. 2015), *S. dunnii* appears in our study as an early diverging taxon in sister position to the four *Salix* species of the *Chamaetia-Vetrix* clade.

To test for whole genome duplication (WGD) events, we examined the distribution of *K*s values between paralogs within the *S. dunnii* genome, together with a dot plot to detect potentially syntenic regions. This revealed a *K*s peak similar to that observed in *Populus*, confirming the previous conclusion that a WGD occurred before the two genera diverged (*K*s around 0.3 in Figure S10) (Tuskan *et al*. 2006). A WGD is also supported by our synteny analysis within *S. dunnii* (Figure 1a, Figure S11). Synteny and collinearity were nevertheless high between *S. dunnii* and *S. purpurea* on all 19 chromosomes, and between the two willow species and *P trichocarpa* for 17 chromosomes (Figure 1c), with a previously known large inter-chromosomal rearrangement between chromosome 1 and chromosome 16 of *Salix* and *Populus* (Figure 1c).

### Identification of the sex determination system

To infer the sex determination system in *S. dunnii*, we sequenced 20 females and 18 males from two wild populations by Illumina short-read sequencing (Table S1). After filtering, we obtained more than 10 Gb of clean reads per sample (Table S14) with average depths of 30 to 40× (Table S15), yielding 4,532,844 high-quality single-nucleotide polymorphisms (SNPs).

A GWAS (genome-wide association study) revealed a small (1,067,232 bp) *S. dunnii* chromosome 7 region, between 6,686,577 and 7,753,809 bp, in which 101 SNPs were significantly associated with sex (Table S16, Figures 2 a&b, Figure S12; Table S16). More than 99% of these candidate sex-linked SNPs are homozygous in all the females, and 63.74% are heterozygous in all the males in our sample (Table S17).

**Fig. 2.**
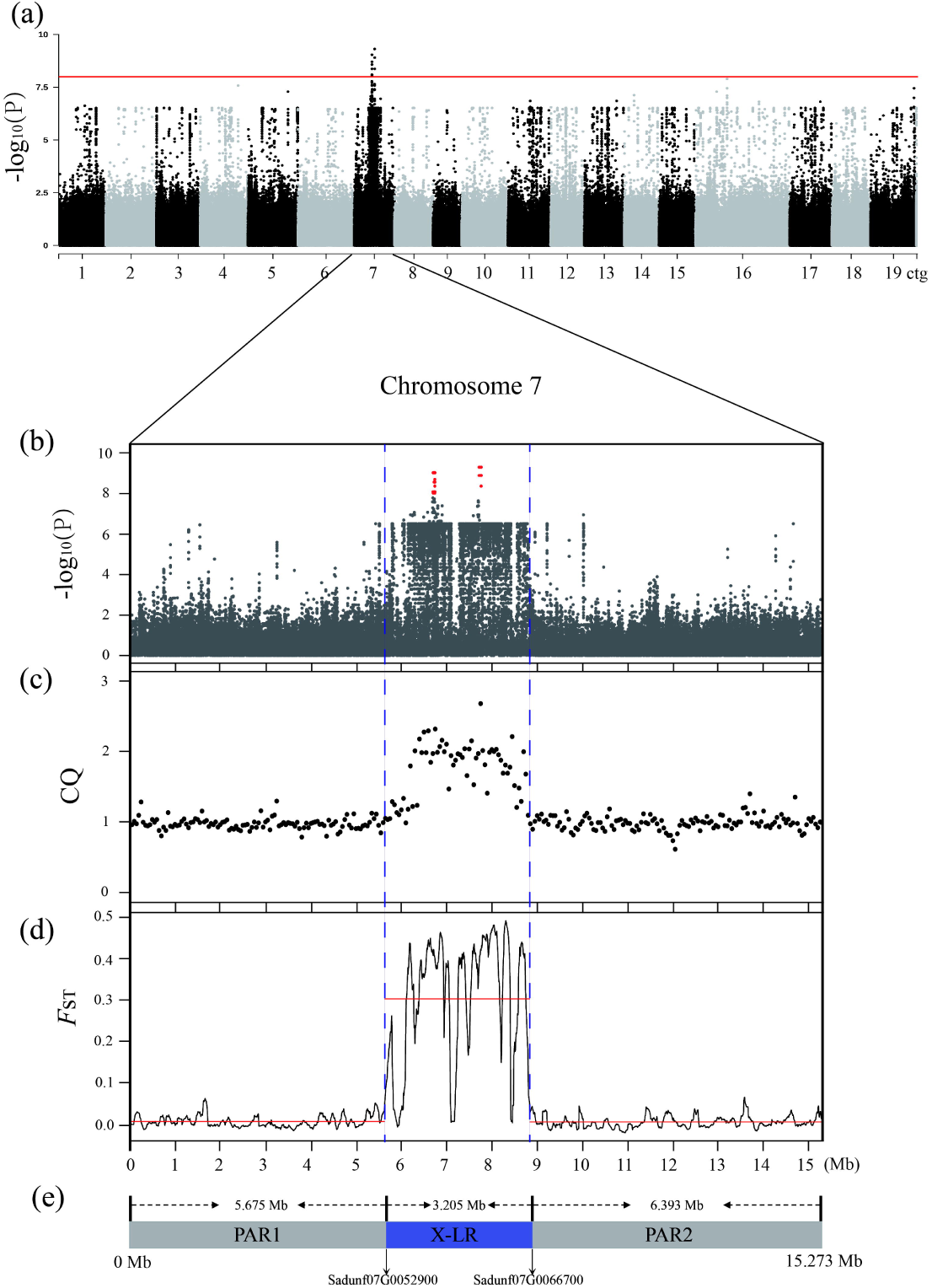
Identification of the sex determination systems of *S. dunnii*. **a**, Results of genome wide association studies (GWAS) between SNPs and sexes in 38 individuals, Q-Q plot for GWAS *P*-values see Figure S12. The Y axis is the negative logarithm of *p* values, and the red line shows the Bonferroni corrected significance level corresponding to α < 0.05. **b**, Manhattan plot for GWAS P-values of all SNPs of chromosome 7. Red dots show significantly sex-associated SNPs. **c**, Chromosome quotients (CQ) in 50 kb non-overlapping window of chromosome 7 (for the rest of the genome, see Figure S13). **d**, *F*_ST_ values between the sexes for 100 kb overlapping windows of chromosome 7 calculated at 5 kb steps. Genome wide *F*_ST_-values are in Figure S14. Red lines represent three significant regions on chromosome 7 suggested by changepoint analysis.

Consistent with our GWAS, the chromosome quotient (CQ) method, with 18 individuals of each sex, detected the same region, and estimated a somewhat larger region, between 6.2 and 8.75 Mb, with CQ > 1.6 (which includes all the candidate sex-linked SNPs), whereas other regions of chromosome 7 and the other 18 chromosomes and contigs have CQ values close to 1 (Figure 2c, Figure S13). These results suggest that *S. dunnii* has a male heterogametic system, with a small completely sex-linked region on chromosome 7. Because these positions are based on sequencing a female, and the species has male heterogamety, we refer to this as the X-linked region (X-LR). We roughly predicted (see Methods) that the chromosome 7 centromere lies between 5.2 and 7.9 Mb, implying that the sex-linked region may be in a low recombination region near this centromere (Figure S1). However, without genetic maps, it is not yet clear whether this species has low recombination near the centromeres of its chromosomes.

Genetic differentiation (estimated as *F*_ST_) between our samples of male and female individuals further confirmed a 3.205 Mb X-LR region in the region detected by the GWAS. Between 5.675 and 8.88 Mb (21% of chromosome 7), changepoint analysis (see Methods) detected *F*_ST_ values significantly higher than those in the flanking regions, as expected for a completely X-linked region (Figure 2, Figure S14). The other 79% of the chromosome forms two pseudo-autosomal regions (PARs) (Figure 2). Linkage disequilibrium (LD) was substantially greater in the putatively fully sex-linked region than in the whole genome (Figure S15).

### Gene content of the fully sex-linked region

We found 124 apparently functional genes in the X-LR (based on intact coding sequences), versus 516 in PAR1 (defined as the chromosome 7 region from position 0 to 5,674,999 bp), and 562 in PAR2 in chromosome 7 (from 8,880,001 to 15,272,728 bp) (Table S9, Table S18). The X-LR gene numbers are only 10.3% of the functional genes on chromosome 7, versus 21% of its physical size, suggesting either a low gene density, or loss of function of genes, either of which could occur in a pericentromeric genome region. We also identified 183 X-linked pseudogenes. Including pseudogenes, X-LR genes form 17% of this chromosome’s gene content, and therefore overall gene density is not much lower than in the PARs. Instead, pseudogenes form a much higher proportion (59%) than in the autosomes (31%), or the PARs (148 and 269 in PAR1 and in PAR2, respectively, or 28% overall, see Table S19, Table S20).

41 genes within the X-linked region had no BLAST hits on chromosome 7 of either *P. trichocarpa* or *S. purpurea* (indicated by green vertical lines in Figure 3c, see also Table S18).

**Fig. 3.**
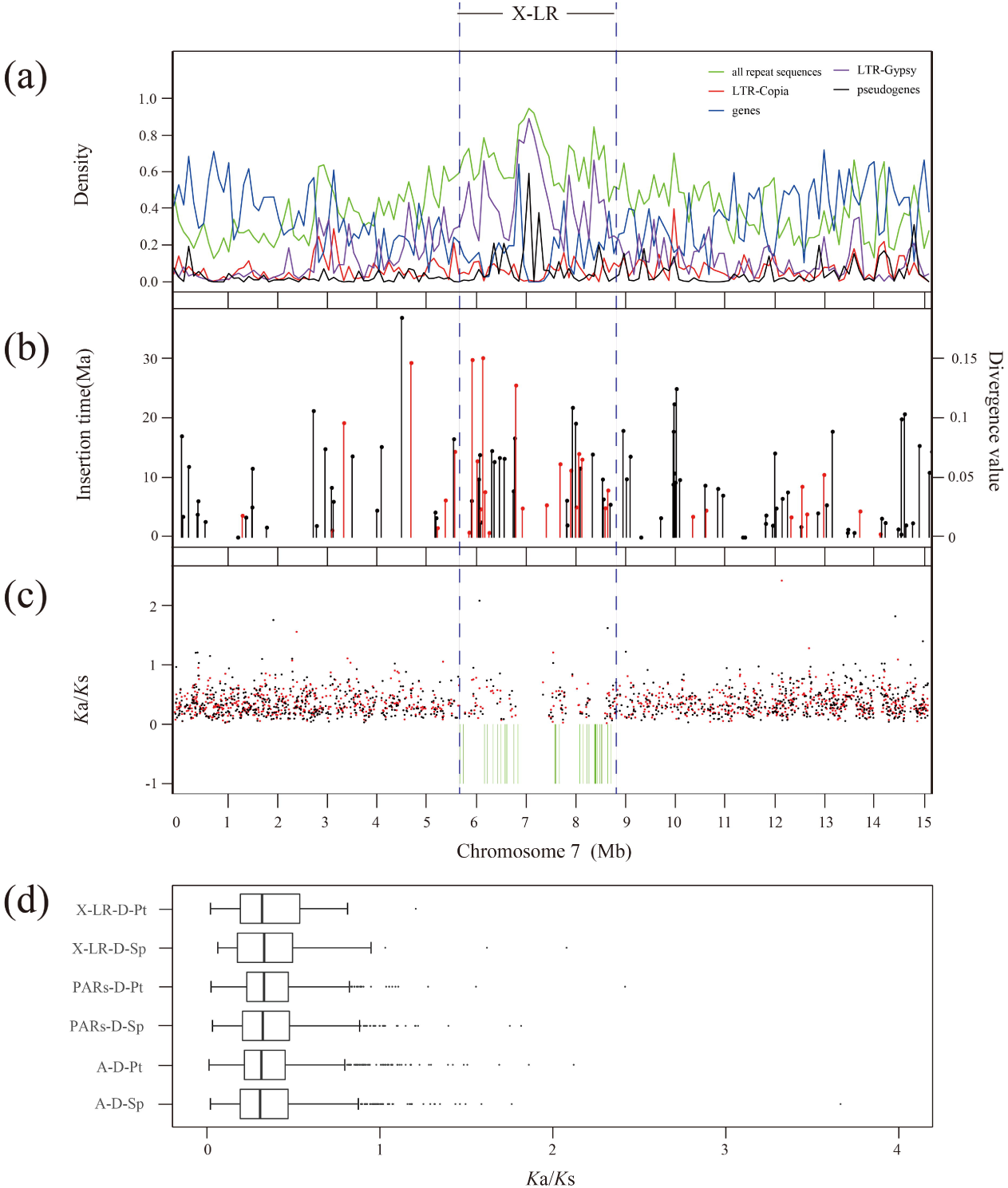
Analysis of *S. dunnii* chromosome 7 genes. **a**, Densities of two transposable element types, LTR-Gypsy (purple line) and LTR-Copia (red line), all repeat sequences (green line), pseudogenes (black line), as well as genes (blue line) in the entire chromosome 7 of *Salix dunnii*. **b**, Estimated insertion times and divergence values of full-length long terminal repeat retrotransposons (LTR-RTs) in chromosome 7 of *S. dunnii*. The red lines represent LTR-Gypsy, and the black lines LTR-Copia elements. **c**, Comparison of *K*a/*K*s ratios between homologous genes in S. *dunnii* and *P. trichocarpa* (red dots), and of *S. dunnii* versus *S. purpurea* (black dots). Green lines indicate locations of *S. dunnii* X-linked genes with no hits in either *S. purpurea* or *P. trichocarpa*. **d**, Comparison of *K*a/*K*s of X-LR, PARs, and autosomal genes (chromosome 6). X-LR -D-Pt and PARs-D-Pt are obtained from the homologous genes of *Salix dunnii* and *Populus trichocarpa*. X-LR-D-Sp and PARs-D-Sp are obtained from chromosome 7 of the homologous genes of chromosome 7of *Salix dunnii* and *Salix purpurea*. A-D-Pt and A-D-Sp are obtained from the homologous genes of chromosome 6 of *Salix dunnii*-*Populus trichocarpa* (1897 homologous pairs) and *Salix dunnii*-*Salix purpurea* (1852 homologous pairs), respectively. The Wilcoxon rank sum test was used to detect the significance difference of different regions of the two datasets. No significant difference (p < 0.05) were detected between the sex-linked region and the autosomes or PARs (Figure S18).

We found a total of eight duplicates or partial duplicates of ARR17-like genes in the *S. dunnii* genome, on chromosomes 1, 3, 8, 10, and 19 (Table S21), but no ortholog or pseudogene copy in the X-LR.

### Molecular evolution of *S. dunnii* X-linked genes

Gene density is lower in the X-LR than the PARs, probably because LTR-Gypsy element density is higher (Figure 3a). Repetitive elements make up 70.58% of the X-LR, versus 40.36% for the PARs, and 40.78% for the 18 autosomes (Table 3). More than half (53.31 %) of the identified intact LTR-Gypsy element of chromosome 7 were from X-LR (Figure 3b, Table S8).

We estimated *K*a, *K*s, and *K*a/*K*s ratios for chromosome 7 genes that are present in both *S. dunnii* and *S. purpurea* (992 ortholog pairs) or *S. dunnii* and *P. trichocarpa* (1017 ortholog pairs). Both *K*a and *K*s values are roughly similar across the whole chromosome (Figure S16 and S17), and the *K*a/*K*s values did not differ significantly between the sex-linked region and the autosomes or PARs (Figure 3d; Figure S18). However, the *K*a and *K*s estimates for PAR genes are both significantly higher than for autosomal genes, suggesting a higher mutation rate (Figure S16 shows the results for divergence from *P. trichocarpa*, and Figure S17 for *S. purpurea*).

**Table 3.**
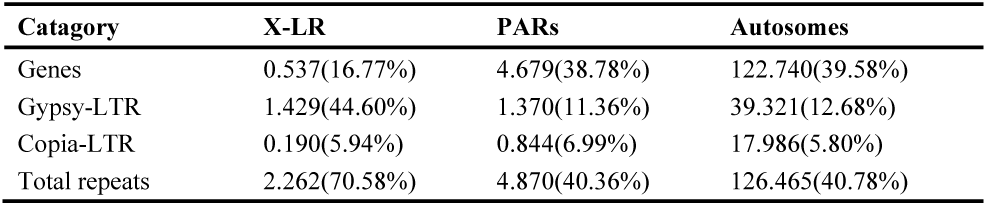
Total size (in Mb) of regions represented by genes and repeat sequences in different regions of the genome (all autosomes were compared with the chromosome 7 X-linked region and its PARs). In parentheses are the proportions of the total lengths of the regions represented by each sequence type.

### Sex-biased gene expression in reproductive and vegetative tissues

After quality control and trimming, more than 80% of our RNAseq reads mapped uniquely to the genome assembly across all samples (Table S22). In both the catkin and leaf datasets, there are significantly more male-than female-biased genes. In catkins, 3,734 genes have sex differences in expression (2,503 male- and 1,231 female-biased genes). Only 43 differentially expressed genes were detected in leaf material (31 male-versus 12 female-biased genes, mostly also differentially expressed in catkins; Figure S19, Table S23). Chromosome 7, as a whole, showed a similar enrichment for genes with male-biased expression (117 male-biased genes, out of 1112 that yielded expression estimates, or 10.52%), but male-biased genes form significantly higher proportions only in the PARs, and not in the X-linked region (Figure 4), which included only 6 male- and 5 female-biased genes, while the other 94 X-LR genes that yielded expression estimates (90%) were unbiased.

**Figure 4.**
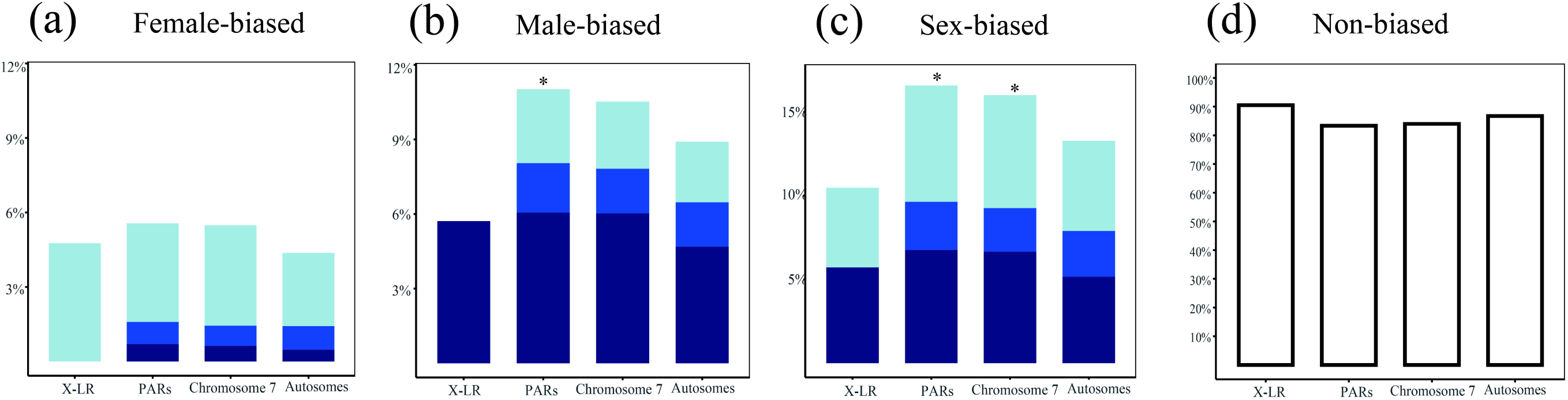
Distribution of sex-biased (| log2FoldChange | > 1, adjusted p value < 0.05) and nonbiased expression genes in catkins. **a**, Female-biased genes. **b**, Male-biased genes. **c**, Sex-biased genes. **d**, Non-biased genes. The percentages of female-biased, male-biased, or non-biased expression genes are shown for different fold change categories (|log2FoldChange|). Light blue bars show values >1, blue indicate values >2, dark blue indicates >3, and open bars are changes less than or equal to twofold. Pearson’s Chi-squared test was used to test the significance difference of sex-based expression genes in different regions (* represent *p* < 0.05).

We divided genes into three groups according to their sex differences in expression, based on the log_2_FoldChange values. All the male biased X-LR genes are in the higher expression category, but higher expression female biased genes are all from the PARs (Figure 4).

## Discussion

### Chromosome-scale genome assembly of *S. dunnii*

The assembled genome size of *S. dunnii* is about 328 Mb (Table 2), similar to other willow genomes (which range from 303.8–357 Mb, Table S24). The base chromosome number for the Salicaceae *sensu lato* family is n=9 or 11, whereas the *Salicaceae sensu stricto* (*s*.*s*.) have a primary chromosome number of n=19 (reviewed in Cronk *et al*. 2015). *Populus* and *Salix* underwent a palaeotetraploidy event that caused a change from n = 11 to n = 22 before the split from closely related genera of this family (e.g. *Idesia*), followed by reduction to n=19 in *Populus* and *Salix* (Darlington & Wylie 1955; Xi *et al*. 2012; Li *et al*. 2019). We confirmed that *Populus* and *Salix* share the same WGD (Figure S10a), and generally show high synteny and collinearity (Figure1c).

### The male heterogametic sex determination system in *Salix dunnii*

The *S. dunnii* sex determination region is located on chromosome 7 (Figure 2), the same chromosome as the only other species previously studied in subgenus *Salix, S. nigra* (Sanderson *et al*. 2020). The size of the X-linked region, 3.205 Mb, is similar to the sizes of Z-linked regions of other willows (Table 1), and they are all longer than any known *Populus* X-linked regions. These data support the view (Yang *et al*. 2020) that sex-determining loci have probably evolved independently within the genus *Salix*, as well as separately in poplars. This is consistent with evidence that, despite dioecy being found in almost all willows, the W-linked sequences of some species began diverging within the genus (Pucholt *et al*. 2017; Zhou *et al*. 2020).

### Gene content evolution in the *S. dunnii* X-linked region

Our synteny analyses and homologous gene identification for the X-LR of our sequenced female support the independent evolution hypothesis (Figure 1c). Many *S. dunnii* X-LR protein-coding genes have homologs on chromosome 7 of *P trichocarpa* and/or *S. purpurea* (Table S18), showing that the region evolved from an ancestral chromosome 7 and was not translocated from another chromosome. However, a third of the protein-coding genes were not found in even the closer outgroup species, *S. purpurea*, whose chromosome 7 is an autosome. These genes appear to have been duplicated into the region from other *S. dunnii* chromosomes, as follows: chromosome 16 (8 genes), 13 (6 genes), 12 (4 genes), 17 (4 genes), 19 (4 genes), and 9 genes from other chromosomes (Table S18). Two of these genes (Sadunf07G0053500 and Sadunf07G0053600) are involved in reproductive processes (respectively resembling the *A. thaliana* genes EMBRYO DEFECTIVE 3003, involved in embryo development and seed dormancy, and CLP-SIMILAR PROTEIN 3, which is involved in flower development), and two others (Sadunf07G0059600 and Sadunf07G0059800) have sex-biased expression (Table S18). However, we cannot conclude that these duplications were selectively advantageous, moving genes with reproductive functions to the X-linked region, as an alternative cannot be excluded (see below).

Given the numerous genes in the *S. dunnii* X-linked region, and the lack of an assembled male genome sequence, no candidate sex determining gene can yet be proposed for this species. In several *Populus* species with male heterogamety, the sex determining gene is a duplicate of a member of the gene family that includes the *ARR17* gene (Xue *et al*. 2020; Müller *et al*. 2020). Such a gene has been suggested to be the sex determining gene of all Salicaceae (Yang *et al*. 2020), based on the finding of *ARR17*-like genes in the W-linked regions of *S. viminalis* and *S. purpurea* (Almeida *et al*. 2020; Zhou *et al*. 2020). No such gene is present in the Z-linked region of *S. viminalis*, consistent with the finding in the *Populus* species that the duplication is carried only in the Y- and not the X-linked region. Our results are consistent with this, as we found no copy or partial duplicate of *ARR17* in the *S. dunnii* X-linked region. However, given that several similar sequences were found in the *S. dunnii* genome, and given the current lack of information about the Y-linked region in this species, it is not clear whether the presence in the *S. purpurea* Z-linked region of a member of this gene family is merely coincidental. Moreover, it seems unlikely that systems with male and female heterogamety could involve the same gene.

As outlined in the Introduction, sex-determining regions often show recombination suppression, leading to genetic degeneration if suppressed recombination persists for enough time. In diploid organisms, only the Y chromosomes are predicted to degenerate, because X chromosomes recombine in the XX females (reviewed in Charlesworth 2015). However, X-as well as Y-linked regions are expected to accumulate repetitive sequences to a greater extent than non-sex-linked genome regions, due to their somewhat lower effective population size, and this has been detected in papaya (Wang *et al*. 2012a). The *S. dunnii* X-LR appears to have done the same, mainly involving accumulation of LTR-Gypsy elements (Table 3; Figures 1a, 3a), and insertions of these elements appear to have occurred after the *Populus* and *Salix* diverged (Figures 1b and 3b) (the two genera diverged about 48–52 Ma (Chen *et al*. 2019)). However, as in papaya, it is not yet clear whether elements have accumulated due to the region having become sex-linked, or because of its location in the chromosome 7 pericentromeric region (Figure S1). The same uncertainty applies to the unexpectedly large numbers of pseudogenes (Table S20) and duplicated genes (Table S18) found in the X-LR compared with other regions of the *S. dunnii* genome.

It was unexpected to find that one third of the genes of *S. dunnii* X-linked genes did not have orthologs on chromosome 7 of either *S. purpurea* or *P. trichocarpa* (Figure 3c, Table S18). These genes appear to have originated by duplications of genes on other *S. dunnii* chromosomes, and some of them may be functional in reproductive or sex-specific processes. However, we did not detect generally elevated *K*a/*K*s ratios in the X-linked region (Figures 3c, 3d, Figure S18), which would be expected for pseudogenes and non-functional gene duplicates, as well for as genes under adaptive changes that might be expected to occur in such a region. Possibly X-linkage evolved too recently to detect such changes, or for many adaptive changes to have occurred, and therefore the picture indicates predominantly purifying selection, similar to the rest of the genome. Overall, the results suggest that transposable element (TE) accumulation may be an earlier change than other evolutionary changes, which is consistent with theoretical predictions that TEs can accumulate very fast (Maside *et al*. 2005). However, it is again unclear whether these changes are due to sex linkage, or to the region being pericentromeric.

### Sex-biased gene expression in reproductive and vegetative tissues

Sex-biased gene expression may evolve in response to conflicting sex-specific selection pressures (Connallon & Knowles 2005). Our expression analysis revealed significantly more genes with male than female biases, mainly confirmed to genes expressed in catkins, and much less in leaf samples (Table S23). This is consistent with observations in other plant species (Muyle, 2019). Male-biased genes were enriched in the *S. dunnii* PARs (Figure 4), but not in the fully X-linked region (Figure 4), unlike the findings in *S. viminalis* (Pucholt *et al*. 2017) where male biased genes appeared to be mildly enriched in the sex-linked region.

## Acknowledgements

This study was financially supported by the National Natural Science Foundation of China (grant No. 31800466) and the Natural Science Foundation of Fujian Province of China (grant No. 2018J01613). We are indebted to Ray Ming, Andrew Brantley Hall, Pedro Almeida, Jia-Hui Chen, Lawrence B. Smart, Zhong-Jian Liu, Xiao-Ru Wang, Wei Zhao, Feng Zhang, Zhen-Yang Liao, Su-Hua Yang, Ya-Chao Wang, Fei-Yi Guo, En-Ze Li, Hui Liu, Shuai Nie, Shan-Shan Zhou, Lian-Fu Chen for their kind help during preparation of our paper.

## Data availability

Sequence data presented in this article can be downloaded from the###.

Accession numbers are listed in ###.

The genome assemblies and annotations are available through Phytozome###.

## Contributions

Li He and Jian-Feng Mao planned and designed the research. Li He, Kai-Hua Jia, Ren-Gang Zhang, Yuan Wang, Tian-Le Shi, Zhi-Chao Li, Si-Wen Zeng, Xin-Jie Cai, Aline Muyle, Ke Yang, and Deborah Charlesworth analyzed data. Li He, Deborah Charlesworth, Kai-Hua Jia, Yuan Wang, Ren-Gang Zhang, Jian-Feng Mao, Natascha Dorothea Wagner, Elvira Hörandl, and Aline Muyle wrote the paper.

## Supplementary information

**Supplementary Note 1:** Ploidy determination

**Supplementary Note 2:** Transcriptome assembly and gene annotation

**Supplementary Note 3:** Comparative phylogeny analysis across willows

## Figure S

**Figure S1** The bottom left part shows genome-wide Hi-C contact interactions of *Salix dunnii*, the upper right part shows the repeat sequences density of *Salix dunnii* genome of each chromosome. The black lines and block show the possible centromeric region of chromosome 7 based on the joint map.

**Figure S2** Flow cytometry histograms of FAFU-HL-1 of *Salix dunnii* (a) and the external standard *S. integra* (b).

**Figure S3** The 17-mer distribution of Illumina PCR-free short-read data. The x-axis shows *K*-mer abundance; the y-axis shows the number of *K*-mer. The solid line represents *Ks* distribution. The dotted red line represents theoretical values.

**Figure S4** Hi-C interaction heatmap of *Salix dunniii* pseudo-chromosome assembly. The resolution used to estimate the interaction strength of each bin is 100 kb.

**Figure S5** Mitochondrial genome of *Salix dunnii*. Genomic features are shown facing outward (positive strand) and inward (negative strand) of the *Salix dunnii* mitochondrial genome represented as a circular molecule. The colour key shows the functional class of the mitochondrial genes. The GC content is represented in the innermost circle.

**Figure S6** Plastid genome of *Salix dunnii*. Genomic features are shown facing outward (positive strand) and inward (negative strand) of the circular *S. dunnii* plastid genome. The colour key shows the functional class of the plastid genes. The GC content is represented in the innermost circle with the inverted repeat (IR) and single copy (SC) regions indicated.

**Figure S7** Insertion time of LTR-RTs (long terminal repeat-retrotransposons) in the genome *Salix dunnii*.

**Figure S8** Proliferation history of different superfamilies of the *Copia* class of LTR-RTs in the *Salix dunnii* genome.

**Figure S9** Proliferation history of different superfamilies of the *Gypsy* class of LTR-RTs in the *Salix dunnii* genome.

**Figure S10** *K*s values distribution for homologous in *Salix brachista, S. dunnii, Spurpurea, S. viminalis, S. suchowenssi*, and *Populus trichocarpa*. (a) five *Salix* species pairs and *P. trichocarpa*; (b) between *P. trichocarpa* and five *Salix* species. *Salix* species and *Populus* species shared the same WGD event with *K*s value about 0.33 and 0.25, respectively. The peaks of divergence of *Populus* and *Salix* is around the *K*s value of 0.14.

**Figure S11** Syntenic dot plot of the self-comparison of *Salix dunnii*.

**Figure S12** Quantile–Quantile (Q–Q) plots of observed and expected GWAS *P*-values. Red dotted line indicates X = Y and blue shading the 95% confidence interval around the expectation of X = Y, that is that allele frequencies and sex are independent.

**Figure S13** Chromosome quotients (CQ) of each 50 kb nonoverlapping window of whole genome of *Salix dunnii*.

**Figure S14** Genome-wide plot of *F*_ST_-values of *Salix dunnii* calculated at 100 kb windows and 5 kb steps.

**Figure S15** Patterns of linkage disequilibrium decay in the whole genome of *Salix dunnii* (a) and in the X-SDR (b). LD is expressed as the squared allele frequency correlation (r^2^) between two sites whose distances apart are indicated on the X-axis.

**Figure S16** Comparing *K*a and *K*s values of *S. dunnii-P. trichocarpa* homologous pairs between the chromosome 7 X-linked region, the two PARs, and autosomes. **a**, *K*a; **b**, *K*s; 990 homologous pairs (excluded 27 homologous pairs with *K*a or *K*s greater than 1) for chromosome 7, and 1846 for autosome (chromosome 6, excluded 51 homologous pairs with *K*a or *K*s greater than 1). **c**, *K*a; **d**, *K*s; 1017 homologous pairs for chromosome 7, and 1897 homologous pairs for autosome. The Wilcoxon rank sum test was used to detect the significant difference (*p* < 0.05). Red lines indicate median of *K*a and *K*s of autosome to make the differences easy to see. **Figure S17** Comparing *K*a and *K*s values of *S. dunnii-S. purpurea* homologous pairs between the chromosome 7 X-linked region, the two PARs, and autosomes. **a**, *K*a; **b**, *K*s; 965 homologous pairs (excluded 25 homologous pairs with *K*a or *K*s greater than 1) for chromosome 7, and 1808 for autosome (chromosome 6, excluded 44 homologous pairs with *K*a or *K*s greater than 1). c, *K*a; d, *K*s; 992 homologous pairs for chromosome 7, and 1852 homologous pairs for autosome. The Wilcoxon rank sum test was used to detect the significant difference (*p* < 0.05). Red lines indicate median of *K*a and *K*s of autosome to make the differences easy to see.

**Figure S18** Comparing *K*a/*K*s ratios between genes of the chromosome 7 X-linked region, the two PARs, and autosomes. **a**, *S. dunnii-P. trichocarpa* homologous pairs. **b**, *S. dunnii-S. purpurea* homologous pairs. The Wilcoxon rank sum test was used to detect the significant difference (*p* < 0.05).

**Figure S19** Venn diagram comparing differential sex-biased expression genes in catkins and leaves.

## Table S

**Table S1** Details of plant materials used in this study.

**Table S2** Assembly statistics of different methods.

**Table S3** Genome datasets used in the paper.

**Table S4** Length statistics of the final reference genome of *Salix dunnii*.

**Table S5** Statistics of the Oxford Nanopore Technologies (ONT) datasets.

**Table S6** Details of DNA-seq and RNA-seq datasets used for assembly and annotation.

**Table S7** Summary of repeat content of the genome of *Salix dunnii*.

**Table S8** The statistics for full-length long terminal repeat-retrotransposons (LTR-RTs) of *Salix dunnii* genome.

**Table S9** Distribution of RNAs on each regions of the genome of *Salix dunnii*.

**Table S10** Statistics of RNAs of the genome of *Salix dunnii*.

**Table S11** Functional annotation of the predicted genes of *Salix dunnii*.

**Table S12** Transcription factor genes from 58 gene families of *Salix dunnii*.

**Table S13** Summary of transcription factor genes of *Salix dunnii*.

**Table S14** Statistics of quality control results of whole genome resequencing datasets.

**Table S15** Summary of mapping results of 38 samples of *Salix dunnii*.

**Table S16** Statistics of significantly sex associated SNPs in the female *Salix dunnii* genome regions.

**Table S17** Statistics of heterozygosity analysis of the 101 sex associated SNPs.

**Table S18** Genes in the X-linked region of *Salix dunnii*.

**Table S19** Pseudogenes on chromosome 7 of *Salix dunnii*.

**Table S20** Comparation of pseudogenes and genes on *Salix dunnii* genome.

**Table S21** Orthologs copies of ARR17 on the whole female genome of *Salix dunnii* searched by tblastn.

**Table S22** Transcriptome data quality control and mapping results.

**Table S23** The numbers of biased gene expression in catkins and leaves.

**Table S24** Statistics of genome size, genes, and sex determination systems of the five willows with assembled genomes.

